# The neurocognitive gains of diagnostic reasoning training using simulated interactive veterinary cases

**DOI:** 10.1101/650499

**Authors:** Maaly Nassar

## Abstract

The present longitudinal study ascertained training-associated transformations in the neural underpinnings of diagnostic reasoning, using a simulation game named “Equine Virtual Farm” (EVF). Twenty participants underwent structural, EVF/task-based and resting-state MRI and diffusion tensor imaging (DTI) before and after completing their training on diagnosing simulated veterinary cases. Comparing playing veterinarian versus seeing a colorful image across training sessions revealed the transition of brain activity from scientific creativity regions pre-training (left middle frontal and temporal gyrus) to insight problem-solving regions post-training (right cerebellum, middle cingulate and medial superior gyrus and left postcentral gyrus). Further, applying linear mixed-effects modelling on graph centrality metrics revealed the central roles of the creative semantic (inferior frontal, middle frontal and angular gyrus and parahippocampus) and reward systems (orbital gyrus, nucleus accumbens and putamen) in driving pre-training diagnostic reasoning; whereas, regions implicated in inductive reasoning (superior temporal and medial postcentral gyrus and parahippocampus) were the main post-training hubs. Lastly, resting-state and DTI analysis revealed post-training effects within the occipitotemporal semantic processing region. Altogether, these results suggest that simulation-based training transforms diagnostic reasoning in novices from regions implicated in creative semantic processing to regions implicated in improvised rule-based problem-solving.

## Introduction

The question of how medical practitioners develop their expert performance has been a subject of research in the fields of cognitive science and informatics over the last 40 years ^1^. Many psychologists and educators have evaluated the validity of diverse cognitive theories, such as semantic network ^2,3^, deliberate practice ^4^, dual-process of thinking ^5,6^ and Bayes’ rule ^7^, in capturing the complexity of clinical reasoning - the cognitive processes involved in disease diagnosis and therapy ^8^. Converging hypothetical evidence suggests that the cognitive underpinnings of professional medical practice pertain to the relational reasoning processes ^9^ involved in: (1) the structural organization of medical concepts within semantic knowledge networks (semantic network theory); (2) the semantic clustering of knowledge into illness scripts (script theory ^10^) through deliberate practice (deliberate practice theory); and (3) the heuristic (pattern recognition ^11^) and analytic (hypothetic-deductive ^12^, Bayesian thinking ^13,14^) clinical problem solving (dual process theory ^15^). Yet, no current studies have directly observed the transformation dynamics of these reasoning processes throughout medical training and practice ^16^.

Rather, most – if not all – of the clinical reasoning research embraced expert-novice study designs that analyzed the intuitive and analytic cognitive processes of clinicians in terms of dual-process theory. These cross-sectional studies have mainly employed think-aloud protocols ^17–19^ and neuroimaging measures ^20–22^ to explore the reasoning strategies used by novice and expert clinicians during answering non- and authorized multiple-choice questions. Think-aloud protocols revealed experts having better organized knowledge structures and involving more cue-related reasoning strategies ^18,19^. Furthermore, analyzing the dual processes of reasoning in experts showed them using both heuristic and analytic reasoning systems simultaneously in solving problems ^17^. Although the think-aloud method is thought to provide insight into the cognitive underpinnings of clinical reasoning, many scholars doubted the validity of verbal reports in reflecting real-time thinking – conscious (analytic) and unconscious (nonanalytic) reasoning processes ^23–25^.

Consequently, studies using functional magnetic resonance imaging (fMRI), have recently explored the neurocognitive correlates of clinical reasoning through dual-process theory ^21,22^. Prefrontal activations showed significant hemispheric differences between students (novices) and practitioners (experts) during analytic reasoning: experts showed significant activities in their right ventrolateral and dorsolateral prefrontal and right parietal cortices; whereas novices demonstrated greater activations in their left ventrolateral prefrontal and left anterior temporal cortex ^22^. According to the authors ^22^, these distinct hemispheric prefrontal activities suggest the involvement of semantic memory and experiential knowledge in novices’ and experts’ analytical reasoning, respectively. Although experience seems to be associated with a left-to-right hemispheric shift in prefrontal activities during analytical reasoning, Durning et al. ^21^ revealed significantly less activation in frontal regions for experts during nonanalytic reasoning. Those identified frontal deactivations were interpreted as an indicator for expert neural efficiency. However, despite the interpretations of the authors ^21,22^, these findings show that experts activate and deactivate their prefrontal regions as a function of clinical case complexity (nonanalytic vs. analytic) rather than as a function of experiential knowledge (novice vs. experts). Thus, it is still unclear whether the development of expertise is associated with concurrent prefrontal neurocognitive transformations. Moreover, whereas expert’s nonanalytic reasoning was associated with greater activations in regions implicated in pattern recognition (inferior occipital gyrus, middle occipital and parahippocampus), rule-based reasoning (cerebellum) and knowledge updating (lateral orbitofrontal cortex), no significant differences were observed in these regions between novices and experts ^21^. This was not entirely surprising as the novices and experts in this study were postgraduate interns and board-certified internists. Thus, both of them seemed to have developed, from their repeated clinical experiences, more abstract knowledge representations, known as illness scripts ^10^, that are believed to allow experts to discern and process meaningful patterns faster ^26^ – another feature of experts’ efficiency. Consequently, these insignificant and discrepant findings raise the question of whether cross-sectional studies are suitable study designs for investigating the neurocognitive transformations that novices go through to become experts.

Indeed, it is extremely challenging to trace and evaluate the neural correlates of reasoning processes throughout clinical training and practice, but current advances in gamification and neuroimaging technologies provide brilliant possibilities to develop simulation games that can accurately replicate real-life scenarios and capture performance, using reliable and objective measures. These gamification technologies have led to a cascade of simulations that attempt to engage medical practitioners in a variety of safe but challenging clinical scenarios that enable repeated practice, trial-and-error learning and problem-solving, and feedback, without jeopardizing patient health. The data gathered from these simulations have clearly emphasized their potential value in capturing expert performance ^27,28^ in medical domains ^29–33^. Nonetheless, none of those simulation-based studies investigated the differences in clinical reasoning across novices and experts. And, no longitudinal studies have used simulators in combination with neural process-tracing measures (neuroimaging) to investigate the cognitive transformations accompanying novice to expert transition. Thus, it is still questionable whether simulation-based trainings can go beyond their entertainment-, engagement- and procedural learning outcomes and establish beneficial neurocognitive foundations for creative clinical reasoning.

The current study presents a game-based paradigm, using a simulation game known as “Equine Virtual Farm” (EVF) ^34^, to investigate the neural foundations of diagnostic reasoning in novices before and after 5-days of training on diagnosing simulated veterinary cases. The present study is the first longitudinal study conducted to ascertain whether the development of expert reasoning is accompanied by parallel transformations in the activity and connectivity of brain networks and whether these transformations will conform to clinical reasoning theories. Based on clinical reasoning theories, it was hypothesized that playing veterinarian without prior experience would involve regions implicated in semantic processing, whereas problem-solving brain regions would be more involved after training.

## Methods

### Subjects

A total of 20 healthy participants (11 females, mean age: 25.65, age range: 20-55) were invited twice to the Center of Cognitive Neuroscience (CCNB) at the Free University Berlin (FUB) to have their pre- and post-training MRI scans. All participants were right handed and had no history of neurological and psychiatric diseases or medications. All sections of the experiment were performed in accordance with the guidelines in the Declaration of Helsinki. The CCNB review board and the ethics committee of FUB approved all procedures. All participants provided written informed consent before MRI scans and were paid 70 € for their participation.

### Experimental design and procedure

Following the designs of cognitive training studies ^35,36^, our study encompassed three phases: (1) the pre-training phase (day 1); (2) the home training phase (days 2-6); and (3) the post-training phase (day 7). The pre-training phase started with inviting participants to have their first MRI scan at CCNB. After receiving information about MRI safety requirements and signing a consent form, participants started their MRI scan with a 5 min structural MRI (sMRI), followed by 20 min task-based functional MRI (fMRI), 10 min eye-opened resting-state MRI (rsMRI) and a final 15 min diffusion tensor imaging (DTI). The home training phase started on the next day and for another 4 consecutive days, during which participants played with EVF (see Supplementary Fig.1) for a minimum of 1 hour per day. To ensure that participants adhered to this training regimen, they were asked to save their work progress and send their saved files (XML files) by email daily. Those XML files included all the laboratory procedures they achieved within the virtual laboratory as well as their diagnostic and therapeutic hypotheses for the horses. Based on their performance, additional tips and reinforcement emails were sent to each participant to keep them on track. After 5 consecutive days of training, each participant was invited again to CCNB to have their last post-training scan and all participants were reimbursed for their participation. Both pre- and post-training scans were similar in terms of acquisition parameters and sequences.

During fMRI, each participant played a veterinarian within EVF for 20 minutes, using a trackball mouse (Fig. 1e). Within those 20 minutes, participants interacted with five horses in the horse yard and examined their physical performance on treadmill and during feeding, drinking and walking. Next, they started collecting and preparing blood samples from horses to perform laboratory tests, followed by writing a laboratory report. And, to reach a final diagnosis, EVF was provided with an office, where participants can read interactive books and scientific papers on a virtual laptop. Because EVF was developed as a stand-alone desktop application, C++ script was implemented into MATLAB to generate scanner-synchronized and accurate presentation of EVF (Fig. 1a, c, d), alternating with static visual stimuli (Vis) (Fig. 1b). Each participant played through a total of 24-time blocks: twelve 80-sec EVF blocks, and twelve 20-sec Vis blocks. To meet the study aims of investigating the neural correlates of analytic reasoning, another version of EVF was developed for the second post-training scan. In that new version, problem-based scenarios (3D animations and positions) were modified and assigned to different horses. These modifications aimed for providing uncertainty conditions, allowing participants to activate their analytic reasoning strategies.

**Figure 1.**
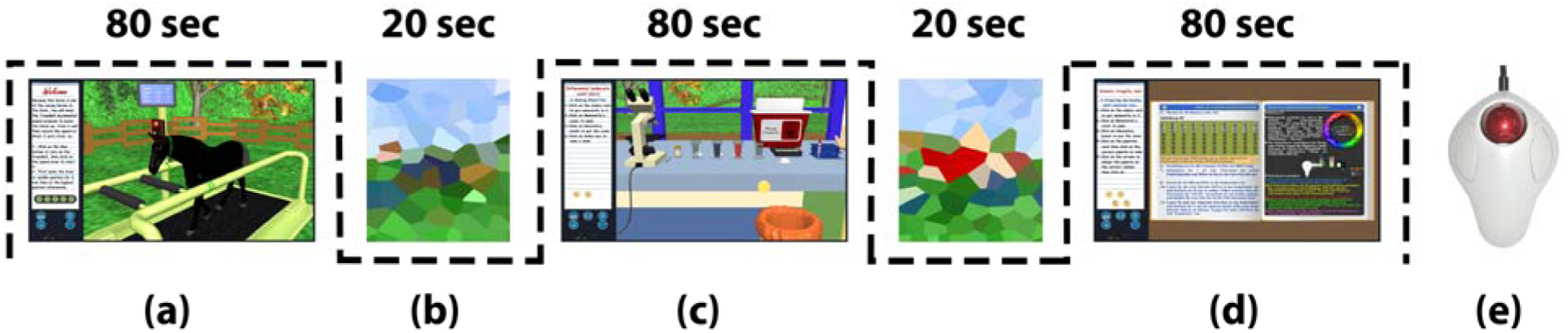
Schematic diagram for the block design of a single fMRI run. Both EVF (80-sec) (a, c, d) and Vis (20-sec) (b) conditions were displayed alternatively for 24-time blocks (12 blocks per condition). During EVF blocks, participants were asked to freely interact with the virtual horses (a) and use the laboratory equipment (c) and books (d) to reach the correct clinical diagnosis. For interaction, subjects used MRI compatible trackball mouse (e).

### Image acquisition

MRI data were collected using a Siemens Vision 3-T Tim Trio scanner (Siemens, Erlangen, Germany) with a standard 12-channel head coil. Anatomical data (sMRI) was acquired using the MPRAGE sequence (TE = 2.52 ms, TR = 1,900 ms, TI = 900 ms, flip angle = 9°, FOV = 256 mm, voxel size = 1 ×1 × 1 mm^3^, 176 sagittal slices). An Echo planar imaging (EPI) sequence (TE = 30 ms, TR = 2,000 ms, flip angle = 70°, voxel size = 3 × 3 × 3 mm^3^, FOV = 192, 37 interleaved axial volumes) was performed for acquiring both task-based (fMRI) and resting-state (rsMRI) data. Finally, Diffusion tensor imaging (DTI) was acquired using an EPI-based single-shot spin-echo diffusion sequence (mz_ep2d_diff_free; TE = 94 ms, TR = 10000 ms, voxel size = 2 × 2 × 2 mm^3^, FOV = 208, phase FOV=100%, 69 transversal slices, Phase Partial Fourier=6/8 flip angle=90°, b-value= 1000s/mm2, bandwidth = 1602 Hz/Px, echo spacing = 0.69 ms, EPI Factor = 104, diffusion directions=61). Distortion correction was implemented using a 3 min point spread function (psf) calibration scan (mz_ep2d_psf) with the same previous diffusion acquisition parameters ^37^. Motion was corrected during reconstruction using a previously acquired reference scan. Total scanning time was ∼1 hour (50 min) encompassing one sMRI, one fMRI, one rsMRI and two DTI scans (mz_ep2d_psf and mz_ep2d_diff_free). Cushions were placed around the head to minimize head movements and stimuli were presented through a mirror mounted on the head coil.

### Image Preprocessing

#### Functional MRI Preprocessing

Images from fMRI, sMRI and rsMRI were preprocessed using SPM12 (Wellcome Trust Center for Neuroimaging, UCL). For each participant, preprocessing started with the realignment of functional images using rigid-motion transform, followed by slice-timing correction. Each structural T1 image was normalized to MNI space, using the unified segmentation and normalization in SPM12, and co-registered to the mean functional image. Then, the estimated parameters of these transformations (i.e. co-registration and normalization) were used to normalize functional and resting-state images (fMRI and rsMRI), which were then smoothed using a Gaussian smoothing kernel (FWHM = 6 mm).

#### DTI preprocessing and tensor fitting

DTI acquisitions were preprocessed using FSL (FMRIB, Oxford, UK, www.fmrib.ox.ac.uk/fsl) Diffusion Toolbox (FDT) (https://fsl.fmrib.ox.ac.uk/fsl/fslwiki/FDT). First, eddy currents and head movements were corrected with the FDT “*eddy_correct”* tool (https://fsl.fmrib.ox.ac.uk/fsl/fslwiki/eddy). Next, non-brain tissue was removed using the FSL brain extraction tool “*bet”* (https://fsl.fmrib.ox.ac.uk/fsl/fslwiki/BET). Finally, fractional anisotropy (FA), mean diffusivity (MD), axial diffusivity (L1) and radial diffusivity (L2, L3) maps were estimated for each participant by fitting a diffusion tensor model at each voxel of the corrected DTI images with the FDT “*dtifit”*.

#### Voxel-based morphometry

A voxel-based morphometry (VBM ^38^) analysis was performed with the Computational Anatomy Toolbox (CAT-12) to investigate the voxel-wise grey matter changes before and after EVF training. CAT-12 default settings were used to segment T1-weighted MRI data into gray matter (GM), white matter, and cerebrospinal fluid, followed by their spatial normalization using the DARTEL template into MNI space. Next, the preprocessed GM data were smoothed using a 6 mm FWHM Gaussian kernel.

### Brain activity Analysis

#### Functional MRI Analysis

For each subject, both fMRI and rsMRI data were modelled using the General Linear Model (GLM). For fMRI data, the GLM included two predictors for EVF and Vis conditions and 6 nuisance regressors for the realignment parameters. In the first-level analysis four contrasts were specified: 1) EVF>baseline [1 0]; 2) Vis>baseline [0 1]; 3) Vis>EVF [-1 1]; 4) EVF>Vis [1 −1]; and t- and beta-maps were generated for each contrast per subject. On the other hand, the GLM of rsMRI included realignment parameters only as six nuisance regressors and no contrasts. Next, between-group analyses were performed on fMRI data using second-level random-effects analysis. Second-level analysis started with performing one sample t-test on EVF>Vis contrasted beta-maps from pre-training (PRE) and post-training (POST) groups, separately. Then, paired t-tests were performed to compare EVF>Vis contrasted beta estimates between PRE and POST groups. Between-group analysis generated two t-maps: 1) pre-vs. post-training (PRE>POST), and 2) post-vs pre-training (POST>PRE). Activation clusters were regarded as significant at P<0.05 FWE-corrected (family-wise error corrected for multiple comparisons) and cluster size k≥10.

### Brain connectivity analysis

#### Functional Connectivity Analysis

Task-based (f-) and resting-state (rs-) connectivity analysis started with extracting the mean time-series of 120 anatomically-defined regions (VOIs) (see Supplementary Table 1), using the “*Volume of Interest”* batch function in SPM12 Utilities. GLM adjusted BOLD responses represented both task-based and resting-state time-series, but task-based time-series were further corrected for EVF>Vis contrast at each voxel (see Supplementary Equations 1-3). Mean time-series were estimated by averaging the voxels composing each anatomical region at each time point, using Neuromorphometrics probabilistic atlas masks (http://www.neuromorphometrics.com). Then, Pearson’s correlation coefficients were computed for each pair of time-series, resulting in the generation of 4 adjacency matrices (2 time-series classes [fMRI and rsMRI] × 2 training sessions) per participant (see Supplementary Fig. 2a). To approach a normal distribution, Fisher’s r-to-z transformation was applied to the correlation coefficients, using Matlab’s *“atanh”* function. (see Supplementary Fig. 2b). Next, the generated adjacency matrices were represented for each subject as undirected graphs G (V,E), where the nodes (V) were the VOIs and the edges (E) were the absolute values of the z-transformed correlation coefficients (weighted edges) ^39^. For each graph, the total number of nodes (V) were 120 and the total number of edges (E) were 1\2 V(V-1) (the upper triangular subset of adjacency matrix; see Supplementary Fig. 2b). Graph centrality metrics, including degree, eigenvector, closeness, betweenness and PageRank ^40^, were then computed for each region, using graph centrality functions in MATLAB2017 (https://de.mathworks.com/help/matlab/ref/graph.centrality.html). To explore the training-associated transformations in brain region centralities, linear mixed-effect models (LME) were fitted for each graph centrality measure, using the *“fitlme”* function in MATLAB2017 (https://de.mathworks.com/help/stats/fitlme.html). For each centrality (C) measure, a linear mixed-effect model (LME) was fitted for each time-series class (i.e. task-based or resting-state), separately, where the centrality metric (C) was the response variable and the interaction between training sessions (T) and VOIs (V) were the fixed effects. Both T and V predictor variables were presented as dummy variables with values indicating their corresponding categorical levels across 20 (subjects) × 117 (VOIs) × 2 (Training) centrality observations. To account for both between- and within subject variance components, LME intercepts were allowed to randomly vary across subjects (S), VOIs grouped by subjects (S:V), or training sessions (T) grouped by VOIs and subjects (S:V:T) (Table 1). Based on previous studies ^41–43^, accounting for these mixed random effects will accurately model the true variability and offer superior statistical power in detecting longitudinal group differences. The resulting LME models were fitted using maximum likelihood estimation (ML) with resting-state, white matter and pre-training set as reference levels (i.e. coefficients set to zero). The random effect structure for each model was independently determined by model comparisons using likelihood ratio test (LRT). LRT compared the observed likelihood ratio (LR) statistic of the compared models with its chi-squared reference distribution and the best fitted models were identified with their appropriate random effects based on “smaller-is-better” Akaike information criterion (AIC) (see Supplementary Table 2). Because the present study aimed for exploring the different central roles of VOIs across training sessions (PRE and POST), significant interaction effects (coefficients) between individual VOIs and training sessions were reported at P<0.05 and visualized using the BrainNet Viewer software (www.nitrc.org/projects/bnv).

**Table 1.** LME models with different random effects. C = Centrality response variable and the fixed-effects for all models are T and V, T = dummy variable representing 2 training groups categories, V = dummy variable representing 117 VOIs categories (4 white-matter regions were set to the same category, hence 117 instead of 120), S = subjects variable, m = 1,2,3…,4800 (centrality observations indices = 20 subjects × 120 VOIs × 2 training groups), k = 1,2 (T categories indices), i =1,2,3 … 117 (V categories indices). S*V= intercept variance across V grouped by S, S*V*T = intercept variance across V grouped by T and S.

|  |  |
| --- | --- |
| $C_{mki} = \beta_0 + \sum_{k=2}^2 \beta_{1k} T_{mk} + \sum_{i=2}^{117} \beta_{2i} V_{mi} + \sum_{k=2}^2 \sum_{i=2}^{117} \beta_{3ki} T_{mk} V_{mi} + \varepsilon_{mki}$ | (1) |
| $C_{mkin} = \beta_0 + \sum_{k=2}^2 \beta_{1k} T_{mk} + \sum_{i=2}^{117} \beta_{2i} V_{mi} + \sum_{k=2}^2 \sum_{i=2}^{117} \beta_{3ki} T_{mk} V_{mi} + \beta_{0n}(S)_n + \varepsilon_{mkin}$ | (1a) |
| $C_{mkin} = \beta_0 + \sum_{k=2}^2 \beta_{1k} T_{mk} + \sum_{i=2}^{117} \beta_{2i} V_{mi} + \sum_{k=2}^2 \sum_{i=2}^{117} \beta_{3ki} T_{mk} V_{mi} + \beta_{0ni}(S * V)_{ni} + \varepsilon_{mkin}$ | (1b) |
| $C_{mkin} = \beta_0 + \sum_{k=2}^2 \beta_{1k} T_{mk} + \sum_{i=2}^{117} \beta_{2i} V_{mi} + \sum_{k=2}^2 \sum_{i=2}^{117} \beta_{3ki} T_{mk} V_{mi} + \beta_{0nik}(S * V * T)_{nik} + \varepsilon_{mkin}$ | (1c) |

#### Structural Connectivity Analysis

Tract-Based Spatial Statistics (TBSS) preprocessing and analysis were performed on the generated DTI parameters maps in FSL. TBSS preprocessing started with the nonlinear alignment of all participants’ FA maps to a standard space (FMRIB58_FA), followed by their affine transformation into MNI152 space. Then, a mean FA image was generated and thinned to create a mean FA skeleton from all subjects’ FA standard-space images. Finally, each participant’s FA, MD, L1, L2 and L3 maps were projected onto the mean FA skeleton after applying a FA threshold of 0.1. The resulting projected maps were then used for TBSS group analyses. To identify significant differences in DTI parameters between PRE and POST groups, voxel-wise paired t-tests were performed on FA, MD, L1, L2 and L3 maps using the nonparametric FSL permutation tool “*randomize”* (http://www.fmrib.ox.ac.uk/fsl/randomise). To correct for multiple comparisons, the contrasts (PRE>□POST, POST□>□PRE) were analyzed with 5000 random permutations using threshold-free cluster enhancement (TFCE) with a significance threshold of P<0.05 FWE-corrected.

### Voxel-based morphometry

To identify the significant differences in grey matter density between PRE and POST groups, volume differences were first assessed for PRE and POST groups using a voxel-wise GLM analysis. Then, paired t-tests were performed to compare GLM beta estimates between PRE and POST groups. Clusters were considered significant at P<0.05 FWE-corrected and cluster size k≥10.

## Results

### Behavioral results

19 of 20 participants successfully completed their functional MRIs and training sessions; one participant successfully completed all other requirements but did not have a pre-training DTI scan. Post-training MRI revealed significant improvement in participants’ laboratory and diagnostic skills. The number of solved cases was significantly higher (p<0.016, t (19) =2.65, SD = 0.76) during the post-training session. However, only 35% of the participants managed to submit an accurate diagnosis for at least one case during their 20-min post-training MRI session. On the other hand, the remaining participants needed more time to gather additional information for corroborating their unresolved premature diagnosis. Most of those remaining participants highlighted after their MRI sessions that despite they knew the correct diagnosis, they needed to test all their hypotheses before submitting their final report. Accordingly, accounting for the reported behavioral results in brain activity and connectivity models was not performed because it was hypothesized that they might lead to faulty neural correlates assumptions.

### Functional activity analysis

Using random-effects analysis, comparing pre-training with post-training sessions (PRE>POST) for EVF>Vis contrast revealed significant activation clusters within the left inferior frontal gyrus (L-IOpFG) and inferior/middle temporal gyrus (L-ITG/MTG). Both clusters overlapped with middle frontal gyrus (MFG) and middle temporal/fusiform gyrus (MTG/FuG), respectively. On the other hand, the reverse contrast (POST>PRE) showed significant cluster activity in the right cerebellum, middle cingulate (R-MCG) and medial superior frontal gyrus (R-MSFG) and left postcentral gyrus (L-POG). (Table 2, Fig. 2).

**Table 2.** Random-effects (between-group) analysis contrasting pre- and post-training (PRE>POST & POST>PRE) for the contrast EVF>Vis. k = cluster size, Z = Z value of peak voxel, L-= left, R-= right, Results are corrected for multiple comparisons (P_FWE-corr_ <0.01). MFG = middle frontal gyrus; IOpFG = inferior frontal gyrus (opercular part); ITG = inferior temporal gyrus; MTG = middle temporal gyrus; FuG = fusiform gyrus; POG = postcentral gyrus; MCG = middle cingulate gyurs; MSFG = medial superior frontal gyrus.

| k | Brain region | Peak MNI |  |  | T | Z |
| --- | --- | --- | --- | --- | --- | --- |
|  |  | x | y | z |  |  |
| PRE>POST |  |  |  |  |  |  |
| 300 | L- MFG/IOpFG | -42 | 11 | 29 | 7.762 | 5.149 |
| 216 | L- ITG/MTG/FuG | -45 | -52 | -13 | 6.916 | 4.831 |
| POST>PRE |  |  |  |  |  |  |
| 106 | R-Cerebellum | 18 | -46 | -22 | 6.393 | 4.615 |
| 245 | L-POG | -36 | -25 | 50 | 6.096 | 4.484 |
| 272 | R-MCG | 3 | -16 | 35 | 5.522 | 4.214 |
| 116 | R-MSFG | 3 | 59 | 26 | 5.206 | 4.054 |

**Figure 2.**
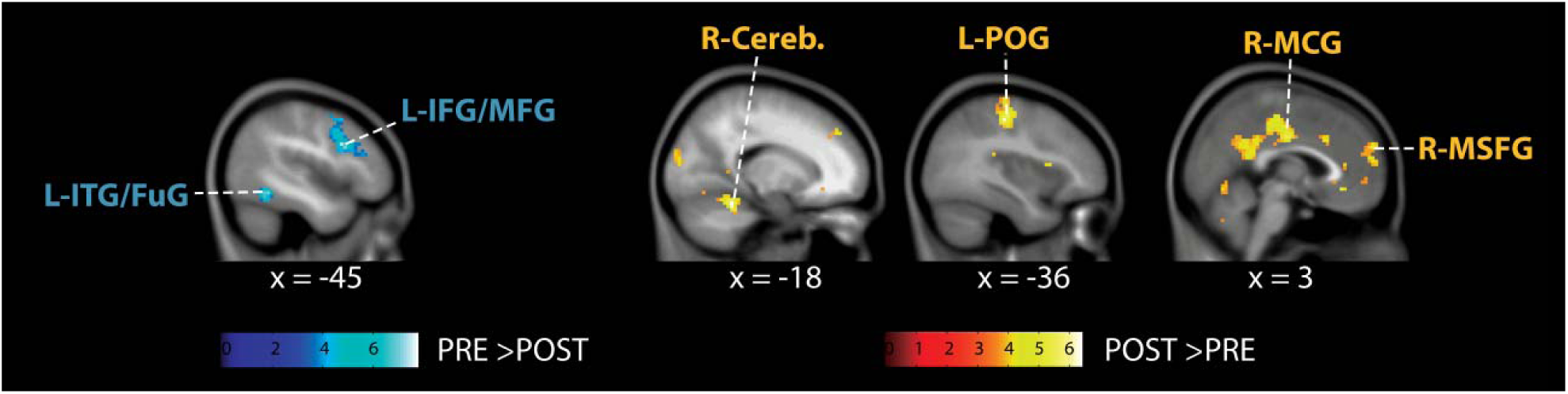
Random-effects (between-group) results for PRE>POST and POST>PRE contrasts, showing significant pre-training activations (blue) in inferior frontal (IFG)/middle frontal (MFG) and inferior temporal (ITG)/fusiform gyrus (FuG) and post-training activation clusters (yellow) in cerebellum (Cereb.), postcentral gyrus (POG), middle cingulate gyrus (MCG) and medial superior frontal cortex (MSFG).

### Functional connectivity analysis

Functional graph analysis started by fitting linear-mixed effects (LME) models for each centrality measure. Model comparisons with different random effects revealed significant improvements and better smaller Akaike Information Criterion (AIC) across centrality measures and MRI groups (see Supplementary Table 2). For both task-based (fMRI) and resting-state (rsMRI) centrality metrics, comparing LME models showed model (1a) fitting significantly better than models (1b) and (1c) for degree and closeness; whereas model (1b) was the best fitted model for eigenvector and PageRank. On the other hand, model (1b) was the best model that explained the variance in betweenness data for fMRI, while model (1c) was the best model for rsMRI. Table 3 and Figure 3A, B focus on the significant interaction between the individual VOIs categories (V) and the training sessions (T) in fMRI and rsMRI. For each centrality measure, regions (V) with significant training (T) interaction coefficients were reported with their related t-values at P<0.05. Within task-based brain network (fMRI), inferior frontal gyrus (IOpFG), angular gyrus (AG), MFG, parahippocampus (PHG) and orbital gyrus (OrFG) showed significant negative post-training effects (negative coefficients) for eigenvector. Similarly, PageRank revealed negative post-training interactions for eigenvector regions as well as superior temporal gyrus (STG) and frontal operculum (FO). In addition, degree models’ coefficients showed only IOpFG with negative post-training effects. Lastly, for betweenness, PHG and medial postcentral gyrus (MPOG) showed positive post-training interactions, whereas the accumbens area, frontal pole (FP), putamen, amygdala and cerebellum showed post-training negative effects (Fig.3a). Alternatively, within resting-state brain network, eigenvector and PageRank LME modelling revealed the fusiform gyurs (FuG), frontal pole (FP) and inferior occipital gyrus (IOG) with positive post-training interactions.

**Table 3.**
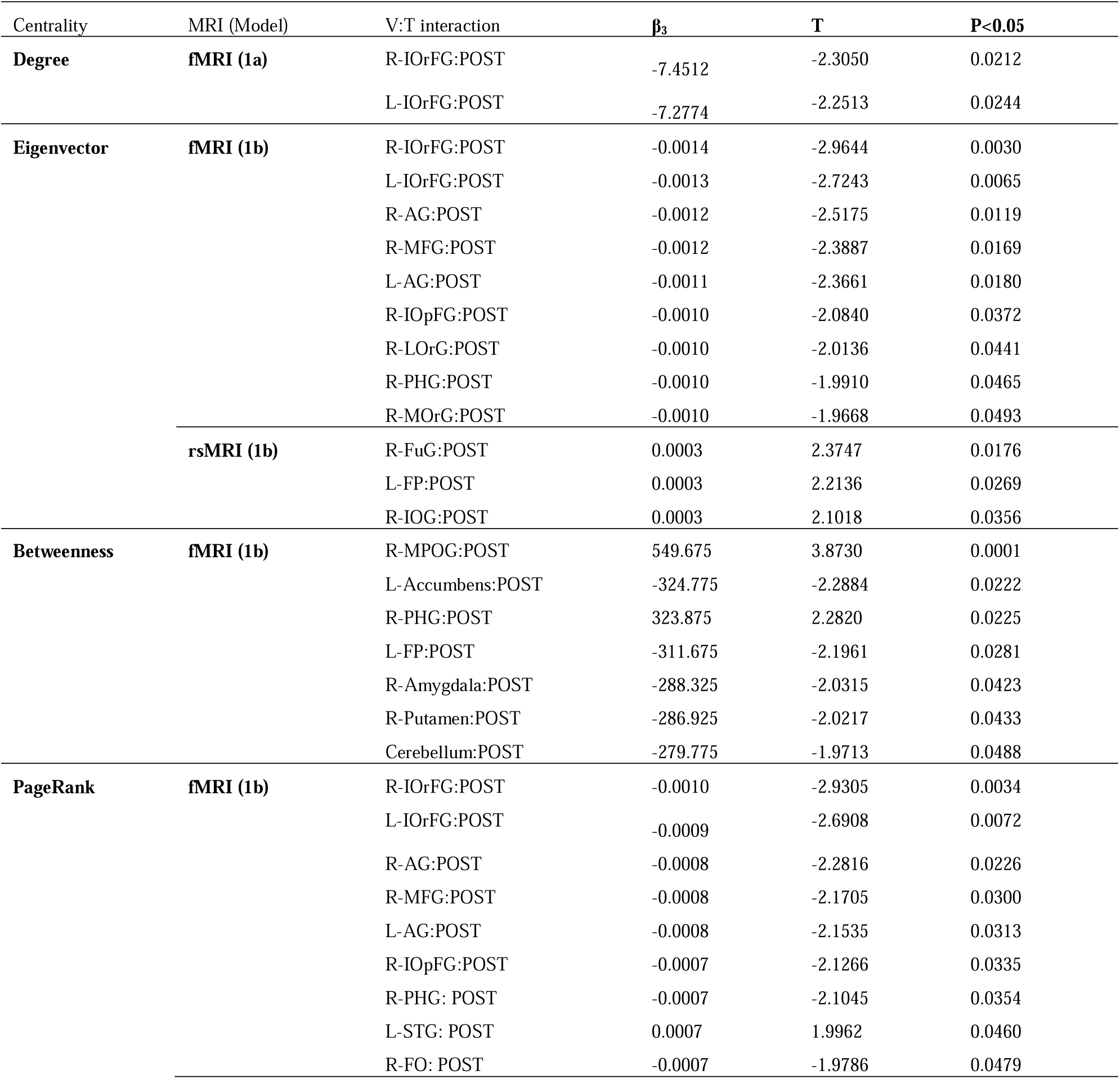
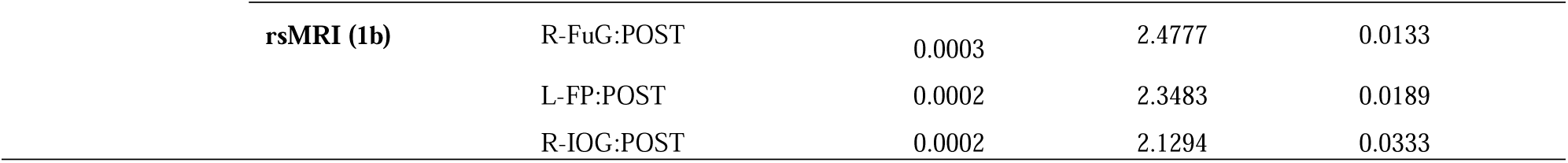
LMEs results for graph centrality measures showing task-based (fMRI) and resting-state (rsMRI) regions (V) with significant (p<0.05, df=4566) post-training (POST) interaction (V:T) coefficients (β3). R-= right; L-= left; IOrFG = inferior frontal gyrus (orbital part); IOpFG = inferior frontal gyrus (opecular part) AG = angular gyrus; MFG = middle frontal gyrus; LOrG = lateral orbital gyurs; PHG = parahippocampus gyrus; MOrG = medial orbital gyurs; FuG = fusiform gyrus; FP = frontal pole; IOG = inferior occipital gyrus; MPOG = medial postcentral gyrus; STG = superior temporal gyrus; FO = frontal operculum.

**Figure 3.**
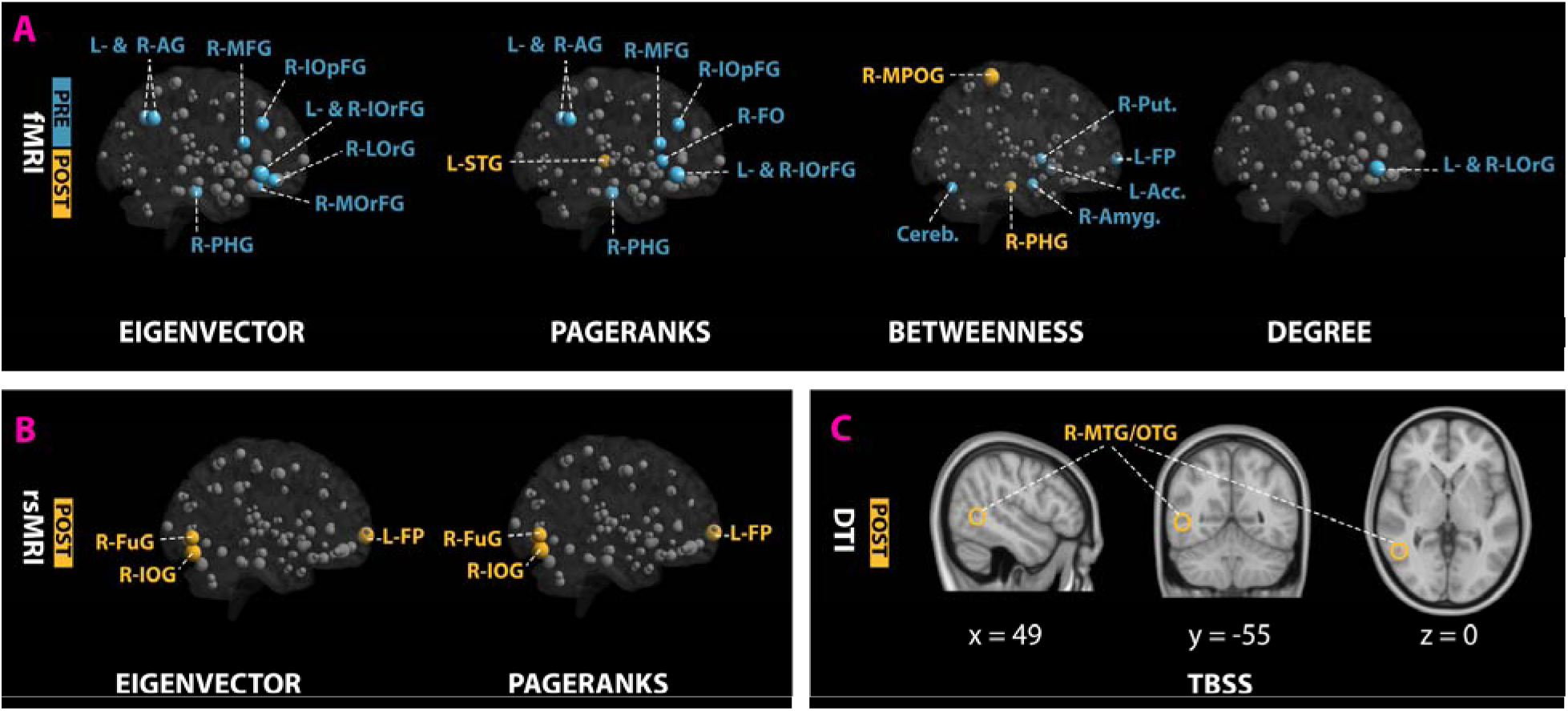
Functional (A,B) and structural (C) connectivity analysis results. Node sizes in fMRI (A) and rsMRI (B) networks indicate the t-values for the V:T interaction coefficients of LME centralities models. Node colors in (A) and (B) indicate regions with insignificant (grey) and significant pre-training (blue) and post-training (yellow) interaction coefficients. For TBSS results (C), yellow brain clusters indicate regions with significant post-training fractional anisotropic values. PRE = pre-training; POST = post-training; R-= right; L-= left; IOrFG = inferior frontal gyrus (orbital part); IOpFG = inferior frontal gyrus (opecular part); AG = angular gyrus; MFG = middle frontal gyrus; LOrG = lateral orbital gyurs; MOrG = medial orbital gyurs;; PHG = parahippocampus gyrus; FO = frontal operculum.; STG = superior temporal gyrus; MPOG = medial postcentral gyrus; Put.=Putamen; Acc.=Accumbens; Amyg.=Amygdala; Cereb. = Cerebellum; FuG = fusiform gyrus; FP = frontal pole; IOG = inferior occipital gyrus; MTG = middle temporal gyrus; OTG=occipitotemporal gyrus.

### Structural connectivity analysis (DTI)

TBSS analyses of multiple diffusion metrics (FA, MD, MO, L1, L2, L3) revealed no significant differences in the mean diffusivity (MD), axial diffusivity (L1) or radial diffusivity (L2, L3). However, fractional anisotropy (FA) showed one significant cluster in right occipitotemporal (ITG/MTG) (t(18)= 7.54, P<0.047 FWE-corrected) (Fig. 3c)

### Voxel-based morphometry (VBM)

T1-weighted MRI data were analyzed using VBM comparing PRE and POST training groups. VBM analysis revealed no significant differences in GM volumes between PRE and POST groups at cluster size k≥10. However, at cluster size k=1, POST training groups showed significant increases in GM within left and right nucleus putamen, caudate and accumbens as well as left thalamus, left Entorhinal area, right anterior insula and right hippocampus.

## Discussion

The present longitudinal study ascertained training-associated transformations in novices’ whole-brain activity and connectivity during diagnostic reasoning, using a simulation game known as Equine Virtual Farm (EVF). Comparing the primary contrast (EVF>Vis) of playing veterinarian (EVF) versus seeing a colorful stationary image (Vis) across training sessions revealed significantly greater activations in the left inferior/middle frontal gyrus (L-MFG/IOpFG) and inferior- or middle temporal/fusiform gyrus (L-ITG/MTG/FuG) during pre-training session (PRE>POST); whereas, the right cerebellum (R-Cerebellum), middle cingulate (R-MCG) and medial superior frontal gyrus (R-MSFG) and left postcentral gyrus (L-POG) showed significantly increased post-training engagement (POST>PRE). Further, graph-based functional connectivity analysis, using LME modelling of centrality metrics, revealed IOpFG, MFG, orbital (OrFG), angular gyrus (AG) and parahippocampus (PHG) with significantly higher connectivity (i.e. eigenvector and PageRank) during pre-training diagnostic reasoning; whereas the superior temporal gyrus (STG) was the only post-training highly connective node. Moreover, LME modelling of betweenness centrality metric showed the central roles of the dopaminergic system (nucleus accumbens and putamen), amygdala and cerebellum in driving pre-training diagnostic reasoning processes; while the PHG and medial postcentral gyrus (MPOG) were the main post-training mediating hubs. On the other hand, resting-state connectivity analysis, using the same task-based LME modelling approach on centrality metrics, revealed higher post-training connectivity for the inferior occipital (IOG) and fusiform gyrus (FuG) and frontal pole (FP). And lastly, structural connectivity analysis using tract-based spatial statistics (TBSS) seems to show diagnostic reasoning training inducing significant structural changes in the occipitotemporal part of middle temporal gyrus (MTG). As hypothesized, these findings suggest the transition of novices’ whole-brain activity and connectivity during diagnostic reasoning from regions implicated in creative semantic processing (MFG/IOpFG, MTG and AG) ^44,45^ before training to regions implicated in improvised rule-based problem solving (cerebellum and POG, MSFG and MCG ^17,46–48^) after training.

The activations of L-MFG/IOpFG have been consistently shown in neuroimaging studies of creative semantic cognition. For instance, Beaty et al. ^45^ have revealed the involvement of both regions in generating unstudied (low-constraint) and novel (high-constraint) semantically related words. In addition, Zhou et al. ^46^ found that searching for numerical relations among conceptual knowledge while solving mathematical problems elicits greater activations in the L-IOpFG, AG, MTG, FuG, PHG, MSFG and posterior cingulate gyrus (PCG). According to Binder et al. ^44^, these seven regions were identified to form the semantic network system. Moreover, on comparing artistic and scientific creativity, Shi et al. ^49^ found a positive correlation between the gray matters of the L-MFG and IOG and scientific creativity, emphasizing the crucial role of semantic reasoning in scientific rather than artistic achievements. Also the significant activities observed in the left MTG and FuG were consistent with a recent meta-analysis study ^50^, where the activities of the left MTG and FuG were attributed to creative ideation during semantic divergent thinking tasks. Thus, the activation pattern observed in the present study within the IOpFG, MFG and FuG suggest that novices relied exclusively on creative semantic processing to explore and reason about the novel relationships among medical concepts, objects (e.g. laboratory equipment) and events (e.g. blood sampling).

Further, modelling graph centrality metrics using LME analysis has extended brain activity findings in the present study and emphasized the central roles of semantic network as well as reward systems in diagnosing novel simulated cases. Specifically, three out of seven semantic network regions ^44^, including IOpFG, AG and PHG, showed higher connectivity (i.e. eigenvector and PageRank) to other brain regions. In addition, the higher connectivity of the orbitofrontal cortex (I-, M- and L-OrFG) suggest that novices relied continuously on monitoring and evaluating the reward values of EVF’s various reinforcers ^51^. Moreover, the evident increases in betweenness centrality metric within the dopaminergic system (nucleus accumbens and putamen), amygdala and cerebellum suggest the incessant contribution of the amygdala, putamen and nucleus accumbens in maintaining the balance between cognitive stability (e.g. inhibiting prepotent responses) and cognitive flexibility (e.g. reward-oriented switching between different options, reversal learning) ^52,53^, while the cerebellum repeatedly simulate proper learning behaviors for providing spontaneous improvisation ^54^. Thus, besides semantic processing, novices seem to engage affective reward processing and improvised creativity in solving novel problems through trial-and-error.

Alternatively, having experience in diagnosing diseases seems to shift brain activity and connectivity from creative semantic processing to insight problem solving. Based on previous neuroimaging studies ^47,48^, the activities in MSFG, POG and cerebellum were found to be associated with a distinct type of mental preparation that leads to successful insight problem solving (spontaneous problem solving, “Aha” experience). Moreover, Crescentini et al. ^55^ found increased activations in STG, cingulate gyrus (CG), cerebellum, putamen and precentral gyrus for rule following during inductive reasoning. These findings seem to connect the observed post-training activations in the cerebellum, POG, MSFG and MCG to rule-based problem solving. Further, the higher connectivity (PageRank) of STG and the increased betweenness of the MPOG emphasize the central roles of semantic priming and rule following in recognizing solutions quickly (insightful problem solving) ^48,56^. Thus, according to the dual process theory ^15^, it is evident that participants engaged the intuitive (associative) and analytical (rule-based) reasoning systems in diagnosing EVF’s swapped simulated cases in EVF new version after training. Also, it is worth noting that in the present study, prefrontal activations shifted with experience from left to right hemispheres. These distinct hemispheric activations were consistent with recent neuroimaging research on clinical decision-making ^22^ that connected the left hemisphere to semantic processing and the right hemisphere to episodic memory retrieval ^57^. These studies explain further the observed engagement of PHG, a region that has been consistently associated with visuospatial processing and episodic memory ^58^. And intriguingly, it is noteworthy that the observed pre- and post-training brain activities were found to be closely similar to the activation pattern implicated in visuo-spatial creative problem solving (MFG, IOpFG, MCG and MSFG) ^59^. This later finding seems to point IOpFG and MFG to creativity and connect MCG and MSFG to problem-solving. However, future studies will be needed to confirm this indication.

Finally, although the post-training changes observed in resting-state and structural connectivity were unexpected, they were consistent with recent neuroimaging research on scientific creativity, which implicated IOG and MTG in scientific semantic processes ^60^ and insight problem solving ^61^. However, a potential limitation might arise from the possibility that these changes might not fully account for EVF training due to the lack of a control group in the present study design.

Taken together, the present study used a novel simulation-based paradigm to study the training-associated transformations in the neural underpinnings of diagnostic reasoning. The present results extend previous neuroimaging studies by clarifying the contribution of semantic processing and insight problem-solving to creative diagnostic reasoning. Moreover, the distinct neural foundations observed in experienced versus inexperienced novices revealed how simulation-based training can shift diagnostic reasoning from creative to rule-based cognitive processes. This neural difference emphasizes the importance of maintaining novelty and challenge within medical training environments for improving the creativity of medical practitioners.

Further, through the LME analysis of functional connectivity, this study showed the contribution of affective (amygdala) and reward-based (nucleus accumbens, putamen, orbital gyrus) processing in driving creative reasoning (middle frontal gyrus and, inferior frontal gyrus) and the engagement of semantic processing (parahippocampus) and insight problem solving (superior temporal) in mediating analytical reasoning. However, whereas resting-state and structural connectivity analysis revealed potential post-training effects within regions implicated in scientific semantic processing, future research should replicate these findings in the presence of control groups.

## Data availability

The datasets generated during and/or analyzed during the current study are available from the corresponding author on reasonable request

## Acknowledgements

This study was funded by the Center for Digital Systems (CeDiS) and supported by the Center of Cognitive Neuroscience Berlin, German Research Foundation (DFG) and the OpenAccess Publication Fund of Freie Universität Berlin. The author would like to thank Prof. Nicolas Apostolopoulos from CeDiS for his technical support, Prof. Felix Blankenburg and Dirk Ostwald from CCNB for their scientific suggestions and revisions and Audrey Hamelers and Christine Ferguson from EMBL-EBI for proof reading the manuscript.

## Additional Information

### Author Contributions

Maaly Nassar designed the study, performed MRI scans, gathered and analyzed the data and wrote the manuscript.

## References

1. Elstein, A. S. Thinking about diagnostic thinking: a 30-year perspective. Adv. Heal. Sci. Educ. 14, 7–18 (2009).

2. Bordage, G. & Lemieux, M. Semantic structures and diagnostic thinking of experts and novices. Acad. Med. 66, S70–2 (1991).

3. Bordage, G., Connell, K. J., Chang, R. W., Gecht, M. R. & Sinacore, J. M. Assessing the semantic content of clinical case presentations: studies of reliability and concurrent validity. Acad. Med. 72, S37–9 (1997).

4. Ericsson, K. A., Krampe, R. T. & Tesch-Römer, C. The role of deliberate practice in the acquisition of expert performance. Psychol. Rev. 100, 363–406 (1993).

5. Norman, G. Dual processing and diagnostic errors. Adv. Heal. Sci. Educ. 14, 37–49 (2009).

6. Croskerry, P. A Universal Model of Diagnostic Reasoning. Acad. Med. 84, 1022–1028 (2009).

7. Westbury, C. F. Bayes’ rule for clinicians: an introduction. Front. Psychol. 1, 192 (2010).

8. Norman, G. Research in clinical reasoning: past history and current trends. Med. Educ. 39, 418–427 (2005).

9. Dumas, D., Torre, D. M. & Durning, S. J. Using Relational Reasoning Strategies to Help Improve Clinical Reasoning Practice. Acad. Med. 93, 709–714 (2018).

10. Custers, E. J. F. M. Thirty years of illness scripts: Theoretical origins and practical applications. Med. Teach. 37, 457–462 (2015).

11. Dumas, D., Alexander, P. A., Baker, L. M., Jablansky, S. & Dunbar, K. N. Relational reasoning in medical education: Patterns in discourse and diagnosis. J. Educ. Psychol. 106, 1021–1035 (2014).

12. Pelaccia, T., Tardif, J., Triby, E. & Charlin, B. An analysis of clinical reasoning through a recent and comprehensive approach: the dual-process theory. Med. Educ. Online 16, (2011).

13. Hall, S. et al. Estimation of post-test probabilities by residents: Bayesian reasoning versus heuristics? Adv. Heal. Sci. Educ. 19, 393–402 (2014).

14. Rottman, B. M., Prochaska, M. T. & Deaño, R. C. Bayesian reasoning in residents’ preliminary diagnoses. Cogn. Res. Princ. Implic. 1, 5 (2016).

15. Evans, J. S. B. T. & Stanovich, K. E. Dual-Process Theories of Higher Cognition. Perspect. Psychol. Sci. 8, 223–241 (2013).

16. Alexander, P. A. Relational thinking and relational reasoning: harnessing the power of patterning. npj Sci. Learn. 1, 16004 (2016).

17. Durning, S. J. et al. Dual processing theory and experts’ reasoning: exploring thinking on national multiple-choice questions. Perspect. Med. Educ. 4, 168–75 (2015).

18. Lesgold, A. et al. Expertise in a complex skill: Diagnosing x-ray pictures. in The nature of expertise. 311–342 (Lawrence Erlbaum Associates, Inc, 1988).

19. Joseph, G.-M. & Patel, V. L. Domain Knowledge and Hypothesis Genenation in Diagnostic Reasoning. Med. Decis. Mak. 10, 31–44 (1990).

20. Durning, S. J. et al. Using functional neuroimaging combined with a think-aloud protocol to explore clinical reasoning expertise in internal medicine. Mil. Med. 177, 72–8 (2012).

21. Durning, S. J. et al. Neural basis of nonanalytical reasoning expertise during clinical evaluation. Brain Behav. 5, e00309 (2015).

22. Hruska, P. et al. Hemispheric activation differences in novice and expert clinicians during clinical decision making. Adv. Heal. Sci. Educ. 21, 921–933 (2016).

23. Russo, J. E., Johnson, E. J. & Stephens, D. L. The validity of verbal protocols. Mem. Cognit. 17, 759–69 (1989).

24. Ericsson, K. A. & Simon, H. A. Verbal reports as data. Psychol. Rev. 87, 215–251 (1980).

25. Fox, M. C., Ericsson, K. A. & Best, R. Do procedures for verbal reporting of thinking have to be reactive? A meta-analysis and recommendations for best reporting methods. Psychol. Bull. 137, 316–344 (2011).

26. Charlin, B., Boshuizen, H. P. A., Custers, E. J. & Feltovich, P. J. Scripts and clinical reasoning. Med. Educ. 41, 1178–1184 (2007).

27. Ericsson, K. A. An expert-performance perspective of research on medical expertise: the study of clinical performance. Med. Educ. 41, 1124–1130 (2007).

28. Ericsson, K. A., Nandagopal, K. & Roring, R. W. Toward a Science of Exceptional Achievement. Ann. N. Y. Acad. Sci. 1172, 199–217 (2009).

29. Iwata, N. et al. Construct validity of the LapVR virtual-reality surgical simulator. Surg. Endosc. 25, 423–428 (2011).

30. Law, B., Atkins, M. S., Kirkpatrick, A. E. & Lomax, A. J. Eye gaze patterns differentiate novice and experts in a virtual laparoscopic surgery training environment. in Proceedings of the Eye tracking research & applications symposium on Eye tracking research & applications - ETRA’2004 41–48 (ACM Press, 2004). doi:10.1145/968363.968370

31. Usón-Gargallo, J. et al. Validation of a Realistic Simulator for Veterinary Gastrointestinal Endoscopy Training. J. Vet. Med. Educ. 41, 209–217 (2014).

32. Williamson, J. A. Construct Validation of a Small-Animal Thoracocentesis Simulator. J. Vet. Med. Educ. 41, 384–389 (2014).

33. Elarbi, M. M., Ragle, C. A., Fransson, B. A. & Farnsworth, K. D. Face, construct, and concurrent validity of a simulation model for laparoscopic ovariectomy in standing horses. J. Am. Vet. Med. Assoc. 253, 92–100 (2018).

34. Nassar, M. Equine virtual farm: A novel interdisciplinary simulation for learning veterinary physiology within clinical context. (Mensch und buch verlag, Berlin, 2011).

35. Brewe, E. et al. Toward a Neurobiological Basis for Understanding Learning in University Modeling Instruction Physics Courses. Front. ICT 5, 10 (2018).

36. Draganski, B. et al. Temporal and Spatial Dynamics of Brain Structure Changes during Extensive Learning. J. Neurosci. 23, 9240–9245 (2006).

37. Zaitsev, M., Hennig, J. & Speck, O. Point spread function mapping with parallel imaging techniques and high acceleration factors: Fast, robust, and flexible method for echo-planar imaging distortion correction. Magn. Reson. Med. 52, 1156–1166 (2004).

38. Ashburner, J. & Friston, K. J. Voxel-Based Morphometry—The Methods. Neuroimage 11, 805–821 (2000).

39. Lohmann, G. et al. Eigenvector Centrality Mapping for Analyzing Connectivity Patterns in fMRI Data of the Human Brain. PLoS One 5, e10232 (2010).

40. Fletcher, J. M. & Wennekers, T. From Structure to Activity: Using Centrality Measures to Predict Neuronal Activity. Int. J. Neural Syst. 28, 1750013 (2018).

41. Bernal-Rusiel, J. L., Greve, D. N., Reuter, M., Fischl, B. & Sabuncu, M. R. Statistical analysis of longitudinal neuroimage data with Linear Mixed Effects models. Neuroimage 66, 249–260 (2013).

42. Chen, G., Taylor, P. A., Shin, Y.-W., Reynolds, R. C. & Cox, R. W. Untangling the relatedness among correlations, Part II: Inter-subject correlation group analysis through linear mixed-effects modeling. Neuroimage 147, 825–840 (2017).

43. Wilson, M. D., Sethi, S., Lein, P. J. & Keil, K. P. Valid statistical approaches for analyzing sholl data: Mixed effects versus simple linear models. J. Neurosci. Methods 279, 33–43 (2017).

44. Binder, J. R., Desai, R. H., Graves, W. W. & Conant, L. L. Where is the semantic system? A critical review and meta-analysis of 120 functional neuroimaging studies. Cereb. Cortex 19, 2767–96 (2009).

45. Beaty, R. E., Christensen, A. P., Benedek, M., Silvia, P. J. & Schacter, D. L. Creative Constraints: Brain Activity and Network Dynamics Underlying Semantic Interference During Idea Production. Neuroimage 148, 189 (2017).

46. Zhou, X. et al. The semantic system is involved in mathematical problem solving. Neuroimage 166, 360–370 (2018).

47. Kounios, J. et al. The prepared mind: Neural activity prior to problem presentation predicts subsequent solution by sudden insight. Psychol. Sci. 17, 882–890 (2006).

48. Tian, F. et al. Neural correlates of mental preparation for successful insight problem solving. Behav. Brain Res. 216, 626–630 (2011).

49. Shi, B., Cao, X., Chen, Q., Zhuang, K. & Qiu, J. Different brain structures associated with artistic and scientific creativity: a voxel-based morphometry study. Sci. Rep. 7, 42911 (2017).

50. Wu, X. et al. A meta-analysis of neuroimaging studies on divergent thinking using activation likelihood estimation. Hum. Brain Mapp. 36, 2703–2718 (2015).

51. Kringelbach, M. L. & Rolls, E. T. The functional neuroanatomy of the human orbitofrontal cortex: Evidence from neuroimaging and neuropsychology. Progress in Neurobiology 72, 341–372 (2004).

52. Jahanshahi, M., Obeso, I., Rothwell, J. C. & Obeso, J. A. A fronto–striato–subthalamic–pallidal network for goal-directed and habitual inhibition. Nat. Rev. Neurosci. 16, 719–732 (2015).

53. Camara, E., Rodriguez-Fornells, A. & Münte, T. F. Functional connectivity of reward processing in the brain. Front. Hum. Neurosci. 2, 19 (2008).

54. Saggar, M. et al. Pictionary-based fMRI paradigm to study the neural correlates of spontaneous improvisation and figural creativity. Sci. Rep. 5, (2015).

55. Crescentini, C. et al. Mechanisms of Rule Acquisition and Rule Following in Inductive Reasoning. J. Neurosci. 31, 7763–7774 (2011).

56. Bowden, E. M. & Jung-Beeman, M. Aha! Insight experience correlates with solution activation in the right hemisphere. Psychon. Bull. Rev. 10, 730–7 (2003).

57. Gazzaniga, M. S. Cerebral specialization and interhemispheric communication: Does the corpus callosum enable the human condition? Brain 123, 1293–1326 (2000).

58. Aminoff, E. M., Kveraga, K. & Bar, M. The role of the parahippocampal cortex in cognition. Trends Cogn. Sci. 17, 379–90 (2013).

59. Kleibeuker, S. W., Koolschijn, P. C. M. P., Jolles, D. D., De Dreu, C. K. W. & Crone, E. A. The neural coding of creative idea generation across adolescence and early adulthood. Front. Hum. Neurosci. 7, 905 (2013).

60. Shi, B., Cao, X., Chen, Q., Zhuang, K. & Qiu, J. Different brain structures associated with artistic and scientific creativity: a voxel-based morphometry study. Sci. Rep. 7, 42911 (2017).

61. Hao, X. et al. Enhancing insight in scientific problem solving by highlighting the functional features of prototypes: An fMRI study. Brain Res. 1534, 46–54 (2013).

