## Supplementary Fig.1, Supplementary Table 1, Supplementary Equations 1-3, Supplementary Fig. 2a, Supplementary Fig. 2b, Supplementary Table 2 for "The neurocognitive gains of diagnostic reasoning training using simulated interactive veterinary cases" (from another version of this preprint: version 1, 2019-05-27)

**Maaly Nassar^1,2,3*^**

^1^ European Molecular Biology Laboratory, European Bioinformatics Institute (EMBL-EBI), Wellcome Trust Genome Campus, Cambridge, UK

^2^ Freie Universität Berlin, Center for Digital Systems, Berlin, 14195, Germany.

^3^ Humboldt-Universität zu Berlin, Berlin School of Mind and Brain, Berlin, 10117, Germany.

^*^Corresponding.

|  | **MNI coordinates** | | |
| --- | --- | --- | --- |
| **Atlas Regions** | **x** | **y** | **z** |
| Cerebellar Vermal Lobules I-V | 0 | -53 | -13 |
| Cerebellar Vermal Lobules VI-VII | 0 | -70 | -23 |
| Cerebellar Vermal Lobules VIII-X | 0 | -61 | -39 |
| Left ACgG anterior cingulate gyrus | -4 | 37 | 8 |
| Left AIns anterior insula | -35 | 11 | -2 |
| Left AOrG anterior orbital gyrus | -24 | 54 | -9 |
| Left Accumbens Area | -9 | 12 | -8 |
| Left Amygdala | -22 | -5 | -20 |
| Left AnG angular gyrus | -43 | -63 | 37 |
| Left Basal Forebrain | -15 | 4 | -15 |
| Left CO central operculum | -48 | -9 | 12 |
| Left Calc calcarine cortex | -9 | -79 | 5 |
| Left Caudate | -13 | 10 | 9 |
| Left Cerebellum Exterior | -24 | -63 | -35 |
| Left Cerebellum White Matter | -18 | -51 | -35 |
| Left Cerebral White Matter | -27 | -17 | 18 |
| Left Cun cuneus | -5 | -80 | 23 |
| Left Ent entorhinal area | -23 | 0 | -28 |
| Left FO frontal operculum | -41 | 18 | 3 |
| Left FRP frontal pole | -10 | 65 | 0 |
| Left FuG fusiform gyrus | -34 | -40 | -21 |
| Left Hippocampus | -25 | -22 | -14 |
| Left IOG inferior occipital gyrus | -40 | -81 | -4 |
| Left ITG inferior temporal gyrus | -48 | -39 | -21 |
| Left LOrG lateral orbital gyrus | -41 | 38 | -15 |
| Left LiG lingual gyrus | -12 | -67 | -6 |
| Left MCgG middle cingulate gyrus | -3 | -2 | 36 |
| Left MFC medial frontal cortex | -5 | 46 | -12 |
| Left MFG middle frontal gyrus | -37 | 30 | 33 |
| Left MOG middle occipital gyrus | -37 | -81 | 22 |
| Left MOrG medial orbital gyrus | -18 | 29 | -18 |
| Left MPoG postcentral gyrus medial segment | -7 | -40 | 64 |
| Left MPrG precentral gyrus medial segment | -6 | -26 | 56 |
| Left MSFG superior frontal gyrus medial segment | -5 | 47 | 25 |
| Left MTG middle temporal gyrus | -57 | -37 | -6 |
| Left OCP occipital pole | -14 | -100 | 6 |
| Left OFuG occipital fusiform gyrus | -27 | -80 | -15 |
| Left OpIFG opercular part of the inferior frontal gyrus | -50 | 15 | 16 |
| Left OrIFG orbital part of the inferior frontal gyrus | -43 | 31 | -8 |
| Left PCgG posterior cingulate gyrus | -5 | -41 | 27 |
| Left PCu precuneus | -6 | -58 | 38 |
| Left PHG parahippocampal gyrus | -21 | -24 | -21 |
| Left PIns posterior insula | -38 | -11 | 3 |
| Left PO parietal operculum | -48 | -31 | 20 |
| Left POrG posterior orbital gyrus | -30 | 21 | -18 |
| Left PP planum polare | -47 | -5 | -5 |
| Left PT planum temporale | -54 | -30 | 13 |
| Left Pallidum | -19 | -3 | -1 |
| Left PoG postcentral gyrus | -40 | -26 | 52 |
| Left PrG precentral gyrus | -39 | -9 | 49 |
| Left Putamen | -25 | 1 | 0 |
| Left SCA subcallosal area | -2 | 17 | -7 |
| Left SFG superior frontal gyrus | -17 | 30 | 48 |
| Left SMC supplementary motor cortex | -5 | 5 | 54 |
| Left SMG supramarginal gyrus | -55 | -39 | 38 |
| Left SOG superior occipital gyrus | -19 | -86 | 31 |
| Left SPL superior parietal lobule | -25 | -57 | 56 |
| Left STG superior temporal gyrus | -58 | -28 | 3 |
| Left TMP temporal pole | -41 | 11 | -30 |
| Left TTG transverse temporal gyrus | -44 | -21 | 9 |
| Left Thalamus Proper | -11 | -18 | 6 |
| Left TrIFG triangular part of the inferior frontal gyrus | -49 | 34 | 5 |
| Right ACgG anterior cingulate gyrus | 5 | 35 | 8 |
| Right AIns anterior insula | 37 | 13 | -2 |
| Right AOrG anterior orbital gyrus | 29 | 44 | -12 |
| Right Accumbens Area | 10 | 13 | -8 |
| Right Amygdala | 22 | -4 | -20 |
| Right AnG angular gyrus | 48 | -57 | 36 |
| Right Basal Forebrain | 18 | 6 | -15 |
| Right CO central operculum | 50 | -6 | 12 |
| Right Calc calcarine cortex | 12 | -76 | 7 |
| Right Caudate | 14 | 10 | 10 |
| Right Cerebellum Exterior | 25 | -62 | -34 |
| Right Cerebellum White Matter | 19 | -51 | -35 |
| Right Cerebral White Matter | 28 | -15 | 18 |
| Right Cun cuneus | 8 | -78 | 24 |
| Right Ent entorhinal area | 24 | 2 | -26 |
| Right FO frontal operculum | 43 | 19 | 3 |
| Right FRP frontal pole | 14 | 66 | 3 |
| Right FuG fusiform gyrus | 35 | -35 | -22 |
| Right Hippocampus | 27 | -20 | -14 |
| Right IOG inferior occipital gyrus | 43 | -78 | -2 |
| Right ITG inferior temporal gyrus | 50 | -38 | -21 |
| Right LOrG lateral orbital gyrus | 41 | 41 | -13 |
| Right LiG lingual gyrus | 14 | -66 | -6 |
| Right MCgG middle cingulate gyrus | 5 | -2 | 37 |
| Right MFC medial frontal cortex | 4 | 44 | -14 |
| Right MFG middle frontal gyrus | 38 | 32 | 32 |
| Right MOG middle occipital gyrus | 41 | -76 | 24 |
| Right MOrG medial orbital gyrus | 20 | 28 | -17 |
| Right MPoG postcentral gyrus medial segment | 7 | -38 | 65 |
| Right MPrG precentral gyrus medial segment | 7 | -25 | 57 |
| Right MSFG superior frontal gyrus medial segment | 6 | 46 | 23 |
| Right MTG middle temporal gyrus | 58 | -34 | -5 |
| Right OCP occipital pole | 19 | -98 | 7 |
| Right OFuG occipital fusiform gyrus | 30 | -77 | -14 |
| Right OpIFG opercular part of the inferior frontal gyrus | 52 | 17 | 17 |
| Right OrIFG orbital part of the inferior frontal gyrus | 45 | 31 | -8 |
| Right PCgG posterior cingulate gyrus | 7 | -40 | 26 |
| Right PCu precuneus | 9 | -57 | 39 |
| Right PHG parahippocampal gyrus | 23 | -22 | -22 |
| Right PIns posterior insula | 39 | -9 | 2 |
| Right PO parietal operculum | 48 | -26 | 22 |
| Right POrG posterior orbital gyrus | 31 | 24 | -17 |
| Right PP planum polare | 48 | -2 | -6 |
| Right PT planum temporale | 55 | -25 | 15 |
| Right Pallidum | 20 | -3 | -2 |
| Right PoG postcentral gyrus | 41 | -23 | 53 |
| Right PrG precentral gyrus | 40 | -6 | 49 |
| Right Putamen | 26 | 3 | 0 |
| Right SFG superior frontal gyrus | 19 | 32 | 47 |
| Right SMC supplementary motor cortex | 6 | 7 | 54 |
| Right SMG supramarginal gyrus | 56 | -33 | 40 |
| Right SOG superior occipital gyrus | 24 | -83 | 31 |
| Right SPL superior parietal lobule | 26 | -54 | 58 |
| Right STG superior temporal gyrus | 59 | -22 | 2 |
| Right TMP temporal pole | 44 | 13 | -28 |
| Right TTG transverse temporal gyrus | 46 | -17 | 8 |
| Right Thalamus Proper | 12 | -18 | 6 |
| Right TrIFG triangular part of the inferior frontal gyrus | 50 | 35 | 4 |

**Table 1.** MNI coordinates of 120 Neuromorphometrics probabilistic atlas masks

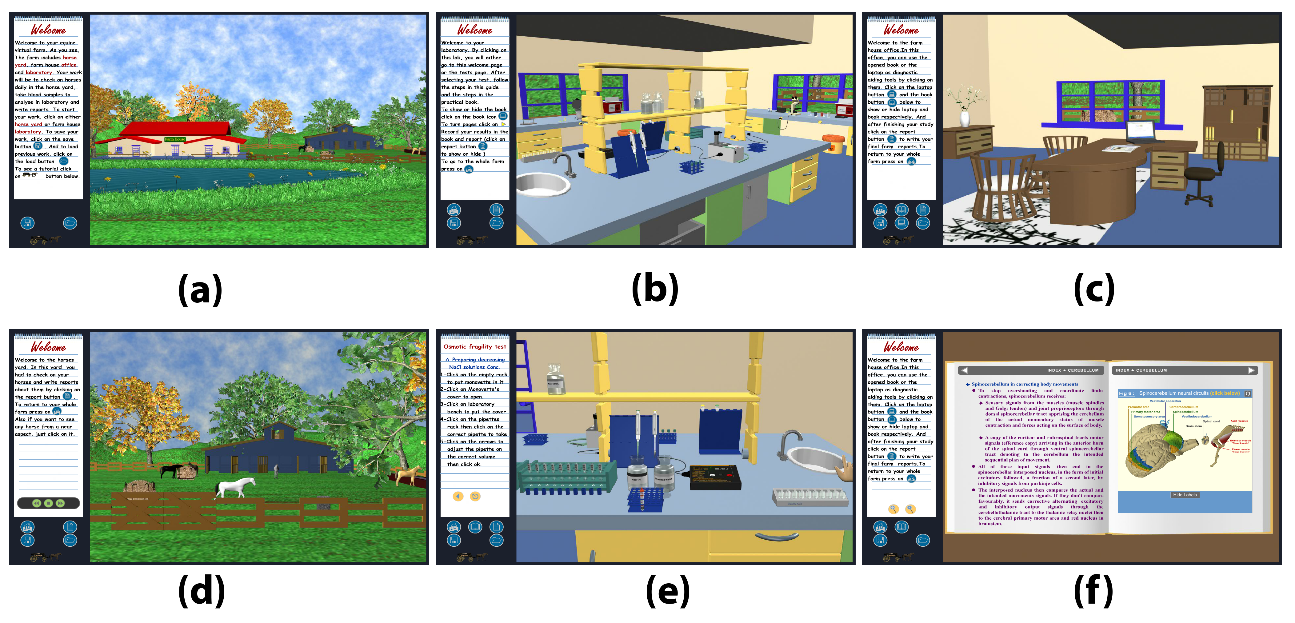

**Figure 1.** Examples of students interactions in the Equine Virtual Farm (a), laboratory (b) , office (c) and horse yard (d): examining the physical performance of horses (d); performing osmotic fragility test on collected blood samples (e); reasoning about the relationships among physiological and clinical concepts (f).

| (a) | (b) |
| --- | --- |
| $R= \begin{matrix} V \\ V_{1} \\ \begin{matrix} V_{2} \\ V_{3} \\ \begin{matrix} \vdots\\ V_{n} \end{matrix} \end{matrix} \end{matrix}\begin{matrix} V_{1} V_{2} V_{3} \cdots V_{n} \\ \left\{ \begin{aligned} r_{11} r_{12} r_{13} \cdots r_{1n} \\ r_{21} r_{22} r_{23} \cdots r_{2n} \\ r_{31} r_{32} r_{33} \cdots r_{3n} \\ \begin{matrix} \vdots& \vdots& \begin{matrix} \vdots& \begin{matrix} \cdots& \vdots\end{matrix} \end{matrix} \end{matrix} \\ r_{n1} r_{n2} r_{n3} \cdots r_{nn} \end{aligned} \right\} \end{matrix}$ | $Z= \begin{matrix} V \\ V_{1} \\ \begin{matrix} V_{2} \\ V_{3} \\ \begin{matrix} \vdots\\ V_{n} \end{matrix} \end{matrix} \end{matrix}\begin{matrix} V_{1} V_{2} V_{3} \cdots V_{n} \\ \left\{ \begin{aligned} z_{11} z_{12} z_{13} \cdots z_{1n} \\ z_{21} z_{22} z_{23} \cdots z_{2n} \\ z_{31} z_{32} z_{33} \cdots z_{3n} \\ \begin{matrix} \vdots& \vdots& \begin{matrix} \vdots& \begin{matrix} \cdots& \vdots\end{matrix} \end{matrix} \end{matrix} \\ z_{n1} z_{n2} z_{n3} \cdots z_{nn} \end{aligned} \right\} \end{matrix}$ |

**Figure 2** Adjacency correlation matrix (R) (a) with its Fisher-transformed counterpart (Z) (b), where n = 120 VOIs (V). As it is a symmetrical matrix, only the upper half of the off-diagonal elements of r-z transformed matrix were considered in undirected graph analysis (red).

| Centrality | MRI Groups | Model Comparison | Model 1AIC | Model 2 AIC | Preferred Model |
| --- | --- | --- | --- | --- | --- |
| **Degree** | **fMRI** | 1a vs. 1b | 35454.6 | 38620.5 | **1a^*^** |
|  |  | 1b vs. 1c | 38620.5 | 39484.7 | 1b |
|  |  | 1a vs. 1c | 35454.6 | 39484.7 | 1a |
|  | **rsMRI** | 1a vs. 1b | 39405.6 | 42854.4 | **1a^*^** |
|  |  | 1b vs. 1c | 42854.4 | 43441.4 | 1b |
|  |  | 1a vs. 1c | 39405.6 | 43441.4 | 1a |
| **Eigenvector** | **fMRI** | 1a vs. 1b | -48165.8 | -48266.6 | **1b^*^** |
|  |  | 1b vs. 1c | -48266.6 | -48165.8 | 1b |
|  |  | 1a vs. 1c | -48165.8 | -48165.8 | - |
|  | **rsMRI** | 1a vs. 1b | -59170.1 | -59854.4 | **1b^*^** |
|  |  | 1b vs. 1c | -59854.4 | -59170.1 | 1b |
|  |  | 1a vs. 1c | -59170.1 | -59170.1 | - |
| **Closeness** | **fMRI** | 1a vs. 1b | -8837.3 | -3681.3 | **1a^*^** |
|  |  | 1b vs. 1c | -3681.3 | -2568.7 | 1b |
|  |  | 1a vs. 1c | -8837.3 | -2568.7 | 1a |
|  | **rsMRI** | 1a vs. 1b | -54508.3 | -50891.2 | **1a^*^** |
|  |  | 1b vs. 1c | -50891.2 | -50557.3 | 1b |
|  |  | 1a vs. 1c | -54508.3 | -50557.3 | 1a |
| **Betweenness** | **fMRI** | 1a vs. 1b | 72456.7 | 72383.1 | **1b^*^** |
|  |  | 1b vs. 1c | 72383.1 | 72456.7 | 1b |
|  |  | 1a vs. 1c | 72456.7 | 72456.7 | - |
|  | **rsMRI** | 1a vs. 1b | 62349.2 | 61405.3 | 1b |
|  |  | 1b vs. 1c | 61405.3 | 59388.6 | **1c^*^** |
|  |  | 1a vs. 1c | 62349.2 | 59388.6 | 1c |
| **PageRank** | **fMRI** | 1a vs. 1b | -51158.0 | -51273.2 | **1b^*^** |
|  |  | 1b vs. 1c | -51273.2 | -51158.0 | 1b |
|  |  | 1a vs. 1c | -51158.0 | -51158.0 | - |
|  | **rsMRI** | 1a vs. 1b | -61015.7 | -61708.3 | **1b^*^** |
|  |  | 1b vs. 1c | -61708.3 | -61015.7 | 1b |
|  |  | 1a vs. 1c | -61015.7 | -61015.7 | - |

**Table 2.** Model comparisons using Akaike Information Criterion (AIC). 1a, 1b, 1c = LME models. * significant AIC at p<0.001.

For all subjects, two classes of time-series were extracted from pre- and post-training GLMs, using the following equations (Eq. 1-3)

$\tilde{Y}=X\tilde{\beta}$ (1)

$Y_{adj}= \tilde{Y}+ \varepsilon$ (2)

$Y_{c}= \tilde{Y}_{c}+ \varepsilon$ (3)

where $\tilde{Y}$ is the estimated predicted response, X is the design matrix with one column per model parameter (i.e. EVF, Vis and motion parameters), $\tilde{\beta}$ are the fitted beta estimates, $\varepsilon$ are the residuals (error terms), $Y_{adj}$ is the adjusted response, $Y_{c}$ is the adjusted response after being corrected for a contrast (EVF>Vis), $\tilde{Y}_{c}$ is the fitted response after being corrected for a contrast (EVF>Vis). Resting-time series was $Y_{adj}$ and task-based time series was $Y_{c}$
